# The discovery of an evolutionarily conserved enhancer within the MYEOV locus suggests an unexpected role for this non-coding region in cancer

**DOI:** 10.1101/2023.09.18.558245

**Authors:** Brigid SA Davidson, Juliana Estefania Arcila-Galvis, Marco Trevisan-Herraz, Aneta Mikulasova, Chris A Brackley, Lisa J Russell, Daniel Rico

**Affiliations:** Biosciences Institute, Newcastle University, Newcastle upon Tyne NE2 4HH, United Kingdom; Translational and Clinical Research Institute, Newcastle University, Newcastle upon Tyne NE2 4HH, United Kingdom; SUPA, School of Physics and Astronomy, University of Edinburgh, Edinburgh EH9 3FD, United Kingdom

## Abstract

The *myeloma overexpressed* gene (*MYEOV*) has been proposed to be a proto-oncogene due to high RNA transcript levels found in multiple cancers, including myeloma, breast, lung, pancreas and esophageal cancer. The presence of an open reading frame (ORF) in humans and other primates suggests protein-coding potential. Yet, we still lack evidence of a functional MYEOV protein. It remains undetermined how *MYEOV* overexpression affects cancerous tissues. In this work, we show that *MYEOV* has originated and may still function as an enhancer, possibly regulating *CCND1*. Firstly, *MYEOV* 3’ end enhancer activity was confirmed in humans using publicly available ATAC-STARR-seq data, performed on B-cell-derived GM12878 cells. We detected enhancer histone marks H3K4me1 and H3K27ac overlapping *MYEOV* in multiple healthy human tissues, which include B cells, liver and lung tissue. The analysis of 3D genome datasets revealed chromatin interactions between the *MYEOV-*3’-enhancer and the proto-oncogene *CCND1*. BLAST searches and multi-sequence alignments results showed that this human enhancer element is conserved from the amphibians/amniotes divergence, with a 273 bp conserved region also found in all mammals, and even in chickens, and it consistently located near the corresponding *CCND1* orthologues. Furthermore, we observed conservation of active enhancer state in the *MYEOV* orthologues of four non-human primates, dog, rat and mouse. When studying this homologous region in mice, where the ORF of *MYEOV* is absent, we not only observed an enhancer chromatin state but also found interactions between the mouse enhancer homolog and *Ccnd1* using 3D-genome interaction data. This is similar to the interaction observed in humans and, interestingly, coincides with CTCF binding sites in both species. Taken together, this suggests that *MYEOV* is a primate-specific gene with a *de novo* ORF that originated at an evolutionarily older enhancer region. This deeply conserved enhancer element is possibly regulating *CCND1* in both humans and mice, opening the possibility of studying *MYEOV* regulatory functions in cancer using non-primate animal models.

## 1. Introduction

*MYEOV* (*myeloma overexpressed*) is a gene believed to be present only in primates (Papamichos et al., 2015). It is proposed to be a proto-oncogene, as its RNA expression has been linked to poorer prognosis in multiple cancers. Cancers showing overexpression of *MYEOV* transcripts include multiple myeloma (Moreaux et al., 2010), breast cancer (Janssen et al., 2002a), esophageal squamous cell carcinomas (Janssen et al., 2002b), gastric cancer (Leyden et al., 2006), colon cancer (Moss et al., 2006), neuroblastoma (Takita et al., 2011) and, more recently, pancreatic ductal adenocarcinoma (PDAC) (Fang et al., 2019; Liang et al., 2020) and non-small cell lung cancer (NSCLC) (Fang et al., 2019). *MYEOV* has also been proposed as a biomarker for prognosis in hepatocellular carcinoma (Deng et al., 2019). However, the exact role that *MYEOV* has in both healthy and cancerous cells remains elusive.

The function of *MYEOV* in healthy cells is currently understudied with competing evidence as to whether it is expressed at very low levels in certain healthy tissues (Liang et al., 2020), with no detectable protein expression in others (Fang et al., 2019). *MYEOV* is believed to have originated with a *de novo* open reading frame (ORF) acquisition during the Catarrhini/Platyrrhini divergence via a mutation leading to the *MYEOV*-313 start codon (Papamichos et al., 2015). Interestingly, *MYEOV*’s protein coding potential is thought to be found only in humans and not in other primates, probably due to a human-specific mutation leading to acquisition of a start codon upstream of the *MYEOV*-255 start codon, extending *MYEOV*’s ORF (Papamichos et al., 2015).

In cancer, it has been shown that the increased RNA expression of *MYEOV* can be caused by genomic rearrangements or duplications of 11q13 (Janssen et al., 2002b; Liang et al., 2020) or by hypomethylation of *MYEOV*’s promoter (Fang et al., 2019; Liang et al., 2020). In PDAC, it was reported that MYEOV protein interacts with the oncogenic transcription factor SOX9 (Liang et al., 2020). MYEOV has been associated with the regulation of microRNAs miR-17-5p and miR-93-5p, possibly by interacting with MYC (Shen et al. 2021). Knockdown of *MYEOV* in pancreatic cell lines suppresses expression of folate metabolic enzymes such as MTHFD2 and restores expression of MYC and mTORC1 repressors (Tange et al. 2023). In NSCLC, it has been proposed to act as a competing endogenous RNA (ceRNA) where it would inhibit the activity of microRNA miR-30c-2-3p (Fang et al., 2019). In fact, a recent paper has also indicated that *MYEOV* might have a role as a ceRNA in PDAC mirroring results seen in NSCLC (Chen et al., 2022).

There is also debate over MYEOV protein function in cancer due to the presence of four upstream translation start sites, believed to render it impossible for *MYEOV* to be translated in human cells (de Almeida et al., 2006). These four upstream AUG sequences located in *MYEOV’*s 5’ untranslated region (UTR) would prevent ribosomal binding via regulation of the ribosomal entry site (Barbosa et al., 2013), abrogating the translation of the *MYEOV* transcript (de Almeida et al., 2006). More recently though a paper reported MYEOV protein expression in PDAC, where they have shown both via immunohistochemistry and western blots that tumour cells exhibit MYEOV protein expression (Liang et al., 2020). This result has not been replicated in NSCLC cell lines where only *MYEOV* RNA expression was observed (Fang et al., 2019). It is possible that *MYEOV* translation might be dependent on cellular states as seen for proinsulin where translational control via upstream AUGs is affected by developmental stages (Hernández-Sánchez et al., 2005).

In our recent study, we explored the chromatin landscape of *MYEOV* in healthy B cells and discovered another possible role for it as an enhancer (Mikulasova et al., 2022). Our previous analyses suggest potential regulatory connections between *MYEOV* and the proto-oncogene *CCND1*. We observed that in the multiple myeloma (MM) cell line U266, the insertion of the immunoglobulin heavy chain (IGH) Eα1 super-enhancer upstream of CCND1 not only changes the chromatin state surrounding *CCND1* but also alters the chromatin configuration of the *MYEOV* gene. We found increased chromosomal accessibility over the *MYEOV* gene body and a H3K4me3 broad domain covering most of the gene (Mikulasova et al., 2022). This is not the first time that *MYEOV* and *CCND1* have been associated together; in fact co-amplification of these two genes are seen in multiple cancers, in particular MM where a 11q13 duplication has occurred (Janssen et al., 2000, 2002b). They have also been linked together in esophageal squamous cell carcinomas where co-amplification of both genes leads to epigenetic silencing of *MYEOV* (Janssen et al., 2000, 2002b) and in primary plasma cell leukaemia where the t(11;14) chromosomal rearrangement leads to *IGH* super-enhancers being juxtaposed next to *MYEOV* and *CCND1* leading to overexpression of both genes (Coccaro et al., 2016). However, the possible role of *MYEOV* as an enhancer and the potential regulatory connections with *CCND1* in healthy cells have not been investigated.

In order to further elucidate *MYEOV*’s possible function in healthy and cancer cells, we have integrated publicly available epigenomics, 3D genome and comparative genomics data to characterise the *MYEOV*-3’-enhancer and elucidate its evolutionary origins. Our data shows that the *MYEOV-*3’-enhancer is a regulatory element that is older than the ORF. The core enhancer sequence is conserved across mammals (even if they lack the ORF), the CCND1 homologue is frequently in synteny, and shows 3D interactions with *MYEOV-*3’-enhancer region in human and mouse cells.

## 2. Material and methods

### 2.1. Quantification of enhancer activity via ATAC-STARR-seq

ATAC-STARR-seq data obtained from GM12878 (Wang et al., 2018a) was used to confirm the enhancer activity of the *MYEOV*-3’-enhancer, measured by the ability of open chromatin regions within the *MYEOV* locus to self-transcribe (the accession number of this dataset, together with the ones corresponding to the other datasets listed below, are listed in **Supplementary table 1**).

### 2.2. Epigenomic datasets

#### 2.2.1. Epigenomic data from human cell types and tissues

ChIP-seq data for H3K4me1, H3K4me3 and H3K27ac histone modifications were retrieved from the NIH Epigenomic Roadmap (Bernstein et al., 2010) to determine the chromatin state surrounding *MYEOV* in human B cells, lung, liver and pancreas (**Supplementary table 1)**.

The entire catalogue (as of December 2021) of human derived H3K27ac ChIP-seq data on the ENCODE database (ENCODE Project Consortium, 2012) was taken in order to determine in which tissues *MYEOV*-3’-enhancer region was active.

#### 2.2.2. Epigenomic data from other species

The chromosomal location of regulatory regions were identified via profiling histone data using ChIP-seq for the five major regulatory histone marks including H3K4me1, H3K4me3, H3K27ac, H3K27me3, and H3K36me3, combined with ATAC-seq in lymphoblastoid cell lines of five primate species (human, chimpanzee, gorilla, orangutan and macaque) (García-Pérez et al., 2021) (**Supplementary Table 1**).

ChIP-seq for H3K4me1, H3K4me3 and H3K27ac histone modifications in CH12 mouse B cell lymphoma cell line, adult liver cells, lung liver cells, and bone marrow, alongside for CTCF from B cells, were retrieved from ENCODE (ENCODE Project Consortium et al., 2020), see **Supplementary Table 3**.

We gathered available ChIP-seq data (H3K4me1, H3K4me3, and H3K27ac) derived from liver cells of five mammalian species (macaque, dog, cat, mouse, rat) from Roller *et al*. (Roller et al., 2021) (**Supplementary Table 1**) with H3K4me3 and H3K27ac ChIP-seq data from the liver of six mammalian species (human, mouse, rat, dog, bovine, and pig) from Villar *et al*., (Villar et al., 2015), and H3K4me3 and H3K27ac ChIP-seq data from the liver of mice (Shen et al., 2012) (**Supplementary Table 1**). For comparison between Roller *et al* and *(Villar et al., 2015)* datasets see **Supplementary Figure 8**.

ChIP-seq datasets for histone marks H3K4me1, H3K4me3 and H3K27ac from chicken liver tissue (Kern et al., 2021) were used to determine chromatin state and sequence conservation of the *MYEOV*-3’-enhancer. **See supplementary Table 1**.

### 2.3. Chromatin conformation data

#### 2.3.1. Chromatin interactions in human cells

Chromatin interaction data were used to determine the possible target of the *MYEOV*-3’-enhancer element, using the data available at the 3D Genome Browser (Wang et al., 2018b). Identification of gene targets of the *MYEOV*-3’-enhancer was obtained using promoter capture Hi-C (PCHi-C) data, for a list of tissues used in this analysis see **Supplementary Table 3**. These cell types were chosen due to *MYEOV* expression being proposed as a prognostic factor for these tissue associated cancers (Moreaux et al., 2010; Fang et al., 2019; Shen et al., 2021). Chromatin Interaction Analysis with Paired-End-Tag sequencing (ChIA-PET) datasets were also obtained, see **Supplementary Table 3**. This was taken alongside CTCF data, **Supplementary Table 3**.

PCHi-C from pre-B cells (Koohy et al., 2018), chromatin capture Hi-C data from mouse embryonic cells (mESC) (Joshi et al., 2015), and PCHi-C data from mESC (Schoenfelder et al., 2015) were used to detect mouse *Ccnd1* promoter interactions with possible target enhancers, see **Supplementary Table 3**. In the pre-B cells only, interactions between 19–22 month mice (“old mice”) were included (Koohy et al., 2018).

#### 2.3.2. Hi-C and HiP-HoP simulations in GM12878 cells

We used the high resolution Hi-C data from Rao *et al*. (Rao et al., 2014) and predicted interactions derived using the HiP-HoP model (Buckle et al., 2018). This modelling approach uses histone modification and DNase I hypersensitivity data to generate a population of 3D chromatin configurations from which simulated Hi-C data can be obtained. The input data and model parameters have been described in detail in our previous study (Rico et al., 2022).

### 2.4. Visualisation of epigenomic and chromatin conformation data

The WashU Epigenome Browser (Li et al., 2022) (http://epigenomegateway.wustl.edu/browser), the 3D Genome Browser (http://3dgenome.fsm.northwestern.edu) (Wang et al., 2018b), Ensembl (v104) website (Martin et al., 2023) and the Integrative Genome Viewer, IGV (Robinson et al., 2011) were used for data visualisation.

### 2.5. Evolutionary analysis of the DNA sequence of *MYEOV*-3’-enhancer

To determine each individual species’ possible enhancer DNA sequence, BLASTN searches were performed using the Ensembl (v104) website (Martin et al., 2023). Briefly, *MYEOV* human enhancer DNA sequence, determined to be GRCh38/hg38 chr11:69293400-69300600 (García-Pérez et al., 2021), was taken from Ensembl and blasted against 17 species (see full list in **Supplementary Table 4**). These searches were under default parameters when the search sensitivity was set to distant homologies, with the exception that the Maximum E-value in which a hit would be reported was set to 1.

Multiple sequence alignment was completed using Mauve with default parameters (Darling et al., 2004). From this alignment, we found the exact region conserved across the different species (**Supplementary Table 4**). Sequences in this region were realigned using the Clustal Omega alignment tool with default parameters (Sievers and Higgins, 2021). This alignment was manually curated using MEGA (Stecher et al., 2020).

## 3. Results

### 3.1 *MYEOV*’s 3’ region has transcriptional enhancer activity in human cells

Chromatin state data from the BLUEPRINT Consortium (Carrillo-de-Santa-Pau et al., 2017) showed that the 3’ end of *MYEOV’s* gene body is encompassed by enhancer chromatin states in healthy B cells (Mikulasova et al., 2022). To investigate *MYEOV*’s possible activity as a transcriptional enhancer, we used ATAC-STARR-seq data generated from the GM12878 lymphoblastoid B cell line (Wang et al., 2018a). ATAC-STARR-seq is based on ATAC-seq isolation of open chromatin regions which are then inserted into a reporter gene within a plasmid, before being transfected into GM12878 cells. If tested open chromatin regions have a regulatory function, they will self-activate, leading to transcription of the reporter gene, measured using RNAseq (Arnold et al., 2013). This experimental approach allows for *in vivo* identification of open chromatin regions which have regulatory functions. Using this method, over 65,000 regulatory regions (active regions) were reported in GM12878 cells, four of which were located in the 3’ UTR region of *MYEOV* (**Figure 1A**). We observed that these regions align with H3K27ac and H3K4me1 peaks and with DNase I hypersensitivity sites in GM12878 cells (**Figure 1A**). Together these observations strongly suggest that this region, located in *MYEOV’s* 3’ UTR, could have an enhancer function.

**Figure 1:**
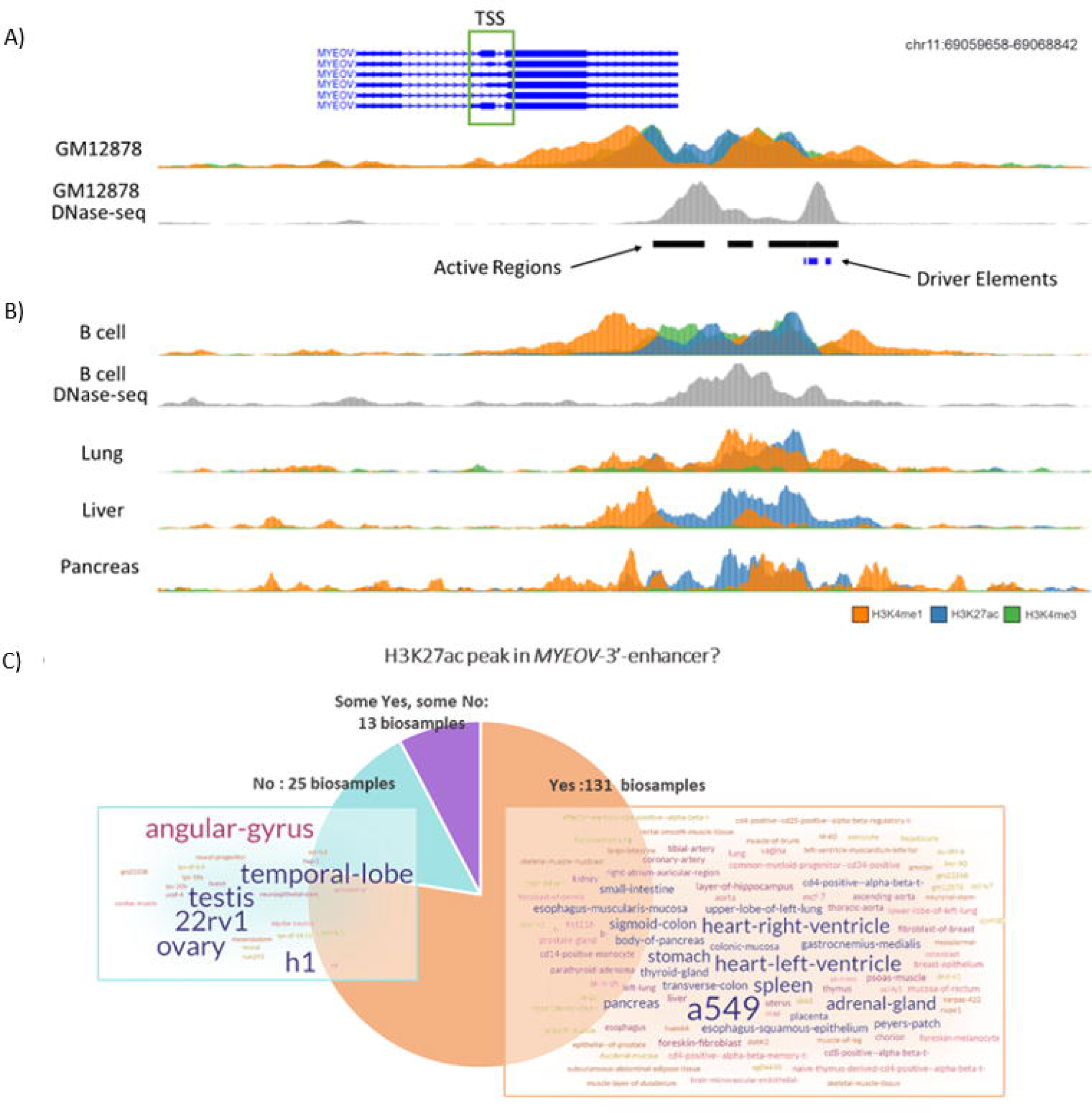
Location of enhancer element within the *MYEOV*’s locus. **A)** ChIP-seq data and DNase-Seq data from lymphoblastoid cell line GM12878 were accessed from the ENCODE data set alongside active regions and driver elements identified using the SHARPR-RE method (Wang et al., 2018a), which predicts sub-regions with high regulatory activity at nucleotide resolution. **B)** ChIP-seq data from B cell, lung, liver and pancreatic tissue together with DNase-seq data in B cells were taken from the ENCODE dataset and visualised using the WashU Epigenome Browser. **C)** Presence of H3K27ac peaks at *MYEOV’*s 3’ UTR region across 169 primary cell samples, tissues and cell lines. NarrowPeak ChIP-seq H3K27ac files were obtained from ENCODE and presence of peaks in chr11:69,297,041-69,298,854 was determined for each tissue. Data shown corresponds to **Supplementary Table 2.** Word clouds show biosample terms specific to groups with and without peaks, darker coloured and larger words represent biosamples with more replicates.

These results therefore strongly indicate that there is a transcriptional enhancer overlapping with *MYEOV*’s 3’ UTR region in GM12878 cells; from now on, we will refer to this region as *MYEOV*-3’-enhancer. In contrast, the histone mark profiles do not indicate an active promoter around the annotated transcription start site (TSS) of *MYEOV* (**Figure 1A**). From this, we next wanted to determine if there are any other cell types in which an equivalent active enhancer is found overlapping *MYEOV*’s 3’ UTR region.

### 3.2 *MYEOV-*3’-enhancer region is active in multiple human cell types and tissues

First we confirmed that the active enhancer chromatin state that was observed in B cells (Mikulasova et al., 2022) using BLUEPRINT data (Carrillo-de-Santa-Pau et al., 2017) could be validated with other datasets. To do this, we used ChIP-seq data for three histone marks (H3K4me1, H3K4me3 and H3K27ac) from the NIH Roadmap (Roadmap Epigenomics Consortium et al., 2015) and ENCODE (ENCODE Project Consortium, 2012) consortia. In the ENCODE datasets we found that the *MYEOV* locus indeed exhibited an enhancer chromatin state in B cells, denoted by the presence of H3K4me1 and H3K27ac (**Figure 1B**). This result is mirrored in data from the NIH Roadmap Consortium, where H3K4me1 and H3K27ac peaks were observed in B cells at the *MYEOV* locus (**Supplementary Figure 1**). This enhancer region was primarily located in the *MYEOV’*s 3’ UTR region, the same location as observed in GM12878 cells (**Figure 1A-B**, **Supplementary Figure 1**).

While the enhancer element is primarily located in *MYEOV*’s 3’ UTR, H3K4me1 is observed across the gene body of *MYEOV* in B cells (**Figure 1B**, **Supplementary Figure 1**). No promoter state (either active or inactive) can be seen at the TSS of *MYEOV.* This is evidenced by the absence of H3K4me3 and H3K27ac peaks in this region, suggesting that *MYEOV* enhancer activity might be independent of *MYEOV* gene transcription in healthy B cells (**Figure 1B**, **Supplementary Figure 1**).

Next, we expanded our search to determine if the active enhancer chromatin state was present in other tissues. We started with three tissues where *MYEOV* expression had previously been linked to cancer including lung (Fang et al., 2019), liver (Deng et al., 2019), and pancreatic tissue (Fang et al., 2019; Shen et al., 2021). In both, the ENCODE (**Figure 1B**) and NIH Epigenomics Roadmap data (**Supplementary Figure 1**), H3K4me1 and H3K27ac signals were observed in *MYEOV’*s 3’ UTR region, highlighting the presence of an active enhancer in this area. Just as in B cells, these cell types showed no active promoter chromatin state at *MYEOV*’s TSS, suggesting that *MYEOV*-3’-enhancer activity is independent of *MYEOV’*s gene promoter activity (**Figure 1B, Supplementary Figure 1**).

With an active enhancer chromatin state found in *MYEOV’s* 3’ UTR in five different cell types, we decided to expand our search using the 182 samples from ENCODE, including primary cells, tissues and cell line samples. We discovered that 131 samples contained H3K27ac peaks within *MYEOV’*s 3’ UTR region (**Figure 1C**, **Supplementary Table 2**), further supporting an enhancer role for this region in humans. Again, the presence of the active chromatin state was associated with tissues in which increased expression is commonly associated with the onset of cancer such as liver, pancreas, colonic mucosa, lung, breast epithelium and stomach smooth muscle (**Supplementary Table 2**). No H3K27ac peaks were observed in 25 tissues, a group that includes brain tissues/primary cells, ovary and testis (**Supplementary Table 2**). To further elucidate this enhancer function and its potential role in cancer development, we aimed to identify possible target genes.

### 3.3. *MYEOV*-3’-enhancer interacts with *CCND1* in human

To determine possible targets of the *MYEOV*-3’-enhancer, we analysed promoter-centred interactions PCHi-C data generated both naive and total B cells (Javierre et al., 2016), where this enhancer element was originally discovered. Here, we observed that in naive B cells, the *MYEOV’*s promoter region interacts with *CCND1* and a gene located downstream, *LTO1* (previously known as *ORAVO1*). In contrast, the *MYEOV*-3’-enhancer interacts only with the region containing the *CCND1* promoter (**Figure 2**). This suggests that in naive B cells, there are two possible targets of this enhancer element. We then investigated whether these interactions were present in other primary cell types. We analysed PCHi-C data from primary tissues and cell lines in which we had not only shown an active enhancer chromatin state in *MYEOV*’s 3’ UTR region (**Figure 1**) but also with documented *MYEOV* overexpression association with cancer (Jung et al. 2019) (see **Supplementary Figure 2**). We observed interactions between *MYEOV* and *CCND1* promoter-containing region in liver, lung, H1-derived mesenchymal stem cells and the GM12878/GM19240 lymphoblastoid cell line (**Supplementary Figure 2**). This further supports our hypothesis that *MYEOV*-3’-enhancer interacts with and could regulate *CCND1*. Unexpectedly, no interactions between *MYEOV* and *CCND1* were seen in pancreatic tissues, despite the presence of histone marks denoting enhancer function surrounding *MYEOV* in this tissue type (**Supplementary Figures 1-3**, **Figure 1A**). Given the observed interaction between *MYEOV* and *LTO1* in naive B cells, we used *LTO1* promoter-containing regions as the viewpoint (**Figure 2**) and only found interactions only with *MYEOV* in lymphoid cells (GM12878/GM1924) and mesenchymal stem cells. When *MYEOV* was used as the viewpoint, an interaction with *LTO1* was observed in lymphoid cells (GM12878/GM1924), mesenchymal stem cells and liver cells (**Supplementary Figures 3-4**). These observations suggest that *CCND1* is the primary target for *MYEOV*-3’-enhancer in the majority of tissues, but it also targets *LTO1*.

**Figure 2:**
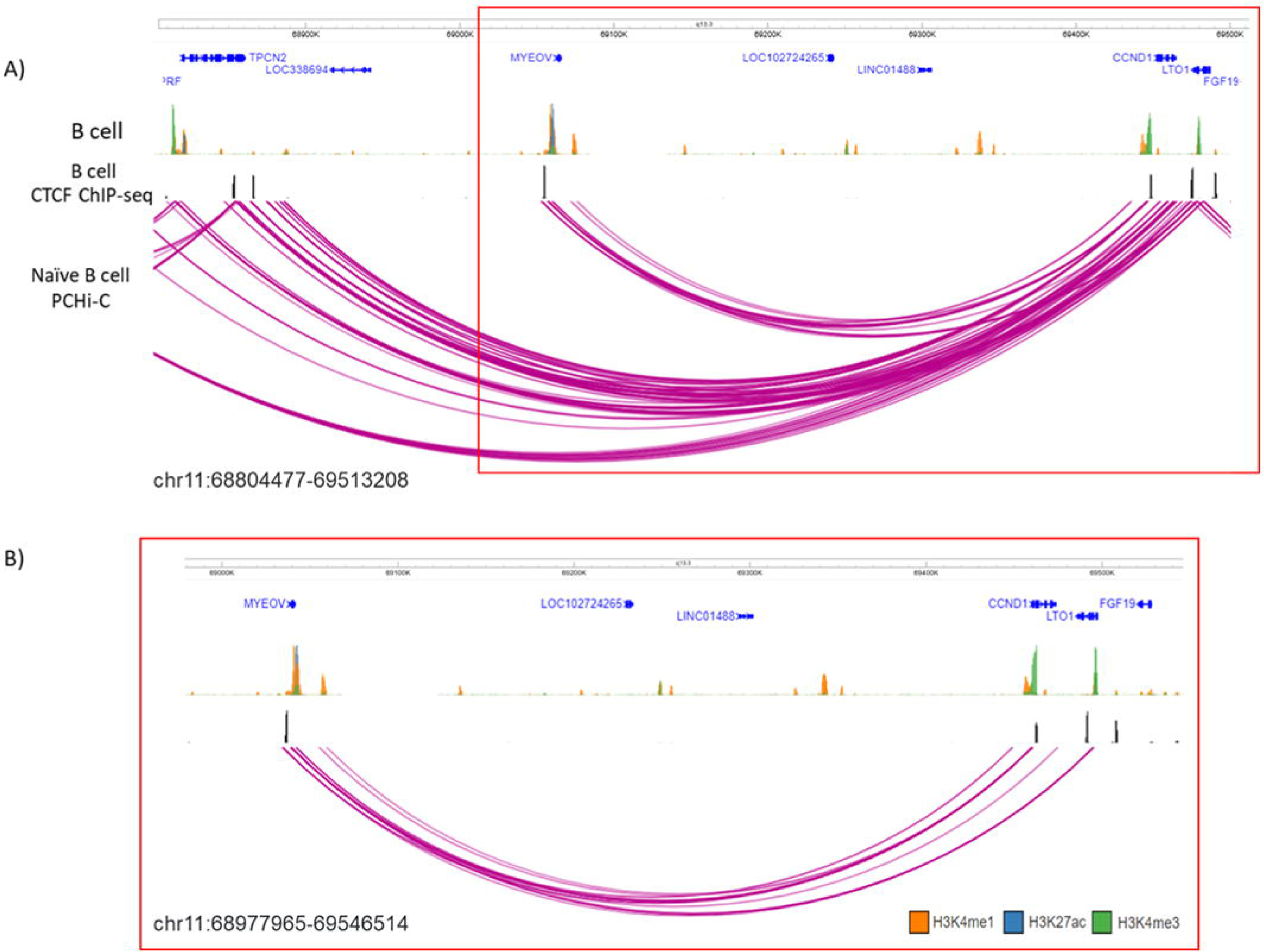
Determination of targets of *MYEOV*’s 3’ UTR enhancer. **A)** ChIP-seq data for H3K4me1 (orange), H3K4me3 (green) and H3K27ac (blue) from B cells appears in top panel, CTCF ChIP-seq data from B cells in middle panel and PCHi-C data from naive B cells in bottom panel, where the purple arcs denote interactions between fragments inside the TAD region containing *CCND1* and *MYOEV*. **B)** Highlight showing that fragments containing *MYEOV* interact with *CCND1*, and *LTO1.* Data is visualised using the WashU Epigenome Browser.

To further strengthen this observation, we analysed ChIA-PET data to determine whether the interaction was mediated via CTCF cohesin loops as we had observed CTCF binding just upstream of *MYEOV* and downstream of *CCND1* in B cells (**Figure 2, 4A**). Here we took CTCF ChIA-PET from the GM12878 cell line (ENCODE Project Consortium, 2012), where interaction between *MYEOV* and *CCND1* had already been detected (**Supplementary 1-3**). We found interactions between the CTCF site located upstream of *MYEOV* and *CCND1*, suggesting these two genes might interact using CTCF cohesin looping (**Figure 3B**). Interestingly, we do not see interactions between *CCND1* and *MYEOV*-3’-enhancer which is downstream of the CTCF site (**Figure 3A-B**). We do see interactions between the *MYEOV*’s CTCF site and another CTCF site located upstream of *MYEOV* which is close to *TPCN2* and possibly highlight the overall structure of this topologically associated domain (TAD) (**Figure 3B**). Another observation is that no interaction is seen between the *MYEOV*’s CTCF site and the CTCF site located close to *LTO1,* which lessens its likelihood as a possible target of *MYEOV*’s 3’ UTR enhancer (**Figure 3B**). To further validate that the interaction between *MYEOV* and *CCND1* is mediated by a CTCF/cohesin complex, we took RAD21 ChIA-PET data (ENCODE Project Consortium, 2012). RAD21 is a subunit of cohesin and therefore highlights interaction mediate via a cohesin complex. This data highlighted the same interaction between the *MYEOV*’s CTCF site and *CCND1* CTCF site (**Figure 3C**). Notably, we did not see any interaction between the *MYEOV* and *TPCN2* CTCF sites (**Figure 3C**). However, this higher-order structure of this TAD could be seen in interactions between the CTCF sites of *CCND1* and *TPNC2* (**Figure 3B-C**). This suggests that the interaction between *MYEOV* and *CCND1* is mediated by a CTCF/cohesin complex not only in GM12878 but also in B cells (**Figure 3A**).

**Figure 3.**
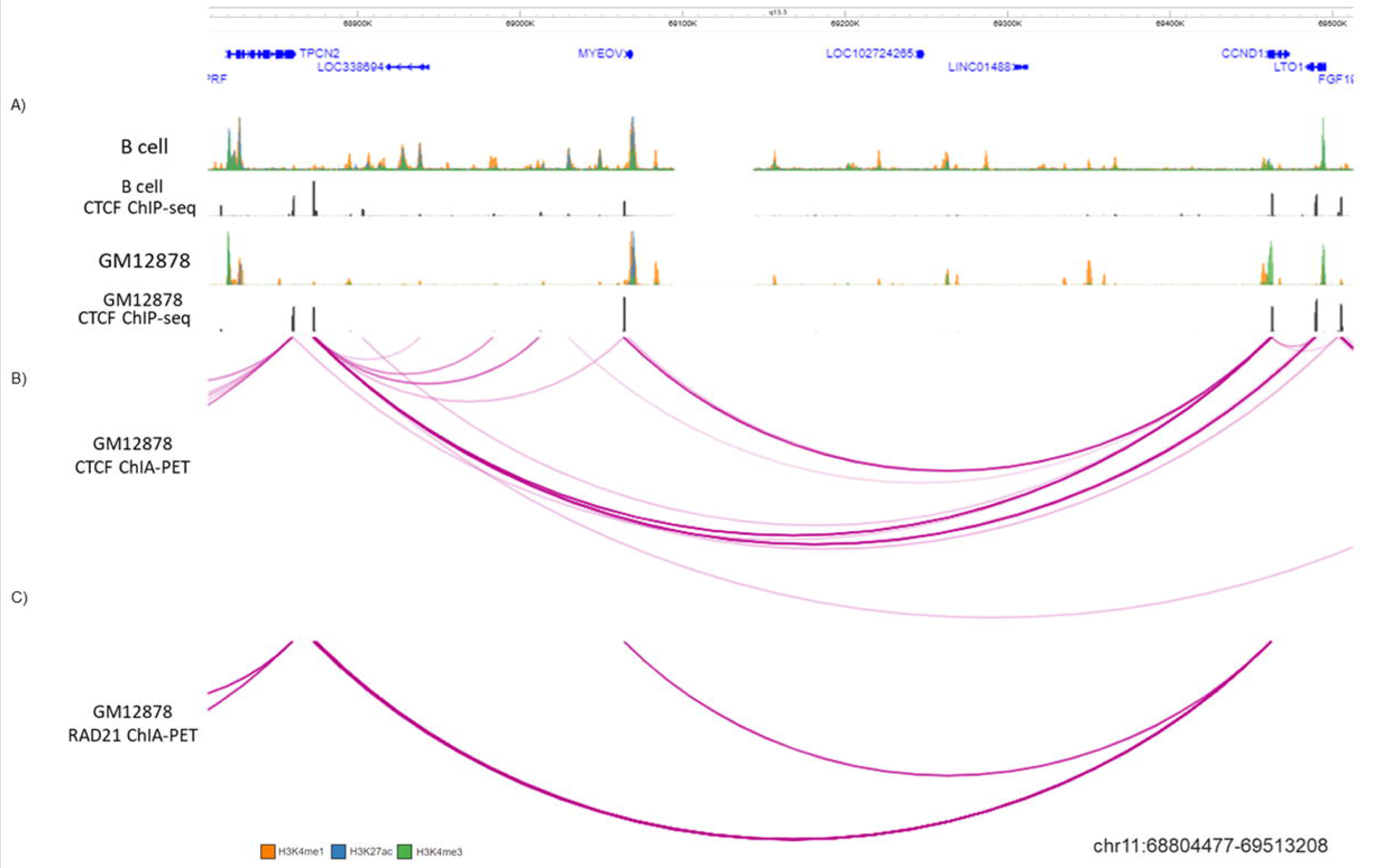
*CCND1* and *MYEOV* interaction is mediated via CTCF cohesion looping. **A)** top first panel shows ChIP-seq data for H3K4me1 (orange), H3K4me3 (green), H3K27ac (blue) second and third panel show CTCF in B cells and histone data in GM12878 cells respectively, **B)** CTCF ChIA-PET data in GM12878 cells, and **C)** RAD-21 ChIA-PET data in GM12878 cells replicates are combined. The purple arcs denote interactions between fragments. Data visualised using the WashU Epigenome Browser.

A more holistic view of chromatin interactions (i.e., not requiring enrichment through promoter capture or immunoprecipitation) can be obtained from Hi-C data. However, Hi-C libraries often lack the complexity to offer sufficient resolution needed to reveal enhancer-promoter interactions. An alternative approach is to use modelling schemes to predict interactions based on other types of epigenomic linear data (Buckle et al., 2018). We recently employed the highly-predictive heteromorphic polymer (HiP-HoP) simulation scheme to study the *CCND1* locus in B cell derived cell lines (Rico et al., 2022). We have previously shown that HiP-HoP provides accurate predictions when compared with high-resolution Capture-C data (Buckle et al., 2018; Rico et al., 2022).

The HiP-HoP model uses the statistical physics of polymers to predict the chromatin configurations in a genomic region using only DNase-seq and ChIP-seq data from three histone modifications and CTCF. The model generates a population of simulated configurations from which Hi-C-like interaction information can be extracted. Three structure-driving mechanisms are included in the model: cohesin-CTCF loop extrusion, chromatin binding protein complexes driving regulatory element interactions and protein condensate formation, and a ‘heteromorphic polymer’ accounting for variation of chromatin fibre properties. **Figure 4** shows Hi-C data at a 10 kbp resolution for GM12878 cells (Rao et al., 2014), alongside our simulated Hi-C data at a 3 kbp resolution for the same cell line (Rico et al., 2022). This predicts that a region around *MYEOV* shows strong interactions with the promoters of *CCND1* and *LTO1*; furthermore, there is a ‘stripe’ of interactions with both of these promoters extending downstream (i.e., 3’) from *MYEOV*. This becomes even more striking when the simulated Hi-C map is normalised against the ‘expected’ interaction level for loci separated by a given genomic distance (bottom row in **Figure 4**). Further interrogation of the simulation reveals that (at least *in silico*) the stripe mainly arises due to loop extrusion by cohesin which becomes halted at the *MYEOV*’s proximal CTCF site; in the model the strong peaks of interaction between DNase I hypersensitive sites within the regulatory element are driven by transcription factor complex binding.

**Figure 4.**
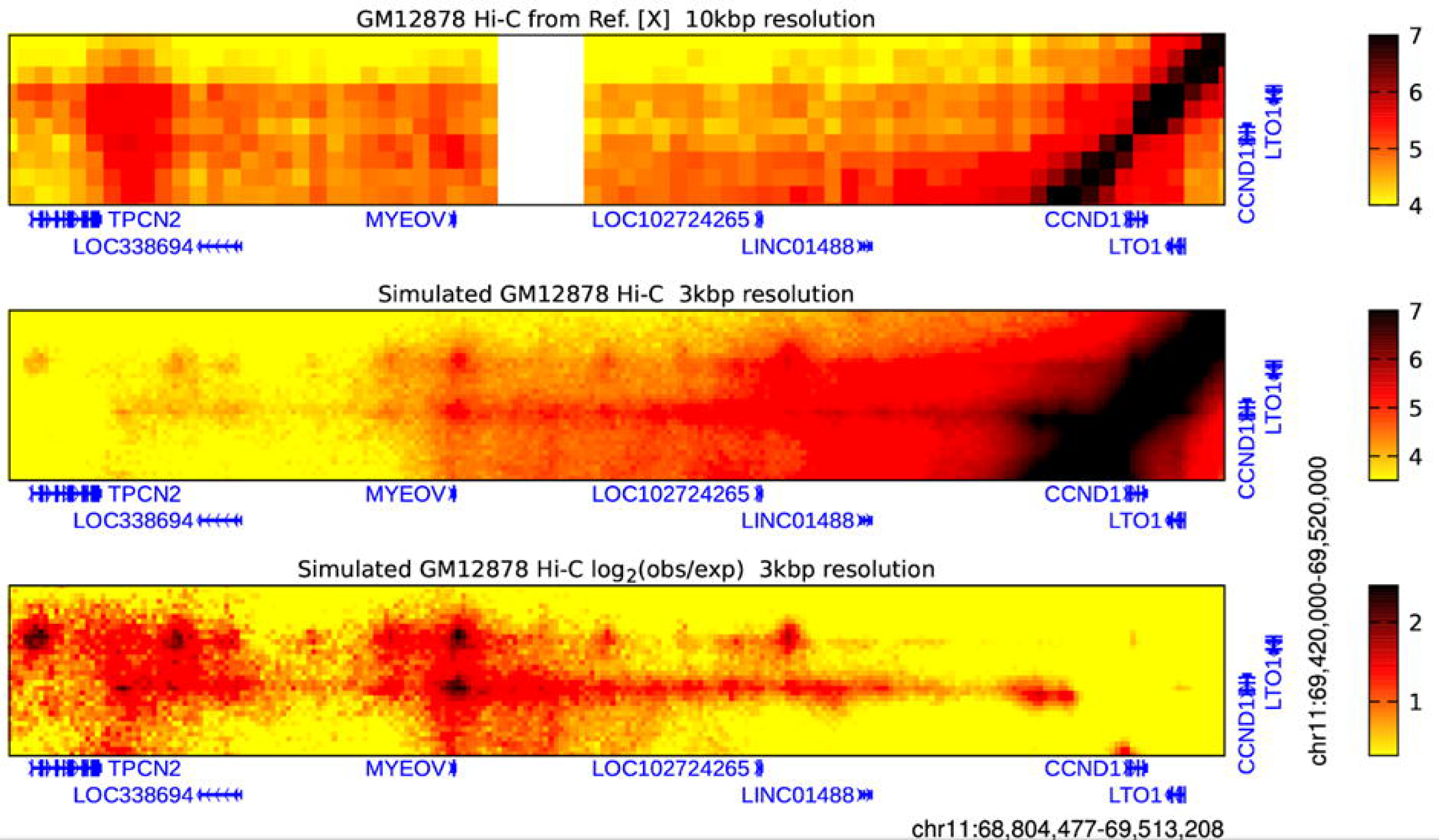
*MYEOV* 3D chromatin interactions evidenced by Hi-C data. Heatmaps showing Hi-C data from GM12878 cells which reveal interactions between a broad region around *MYEOV* and a smaller region enclosing just *CCND1* and *LTO1*, as indicated. Top: Hi-C data are binned at 10 kbp resolution (Rao et al., 2014). Colour scale units are log-normalised interaction counts; the darker the colour, the more frequently two genomic loci were found proximal within a population of cells. An unmappable repetitive sequence region is shown in white. Middle: interactions at higher resolutions can be predicted using the HiP-HoP polymer simulation scheme (Rico et al., 2022). Interactions for the same region as above are shown binned at 3 kbp resolution; colour scale units are log simulated interaction counts. Here, the darker the colour, the more frequently two genomic loci were found proximal within a population of simulated chromatin configurations. Bottom: simulated Hi-C is plotted as log2 observed over expected interaction counts. The ‘expected’ interactions are obtained by calculating the mean number of interactions observed for a given genomic separation.

The experimental data and the results from the simulations strongly suggest that the human *MYEOV-*3’-enhancer functions as a regulatory element that interacts with the proto-oncogene *CCND1*. The ORF of *MYEOV* is found only in primates (Papamichos et al., 2015) but we do not know if the *MYEOV-*3’-enhancer originated around the same evolutionary time. Therefore, we decided to investigate the evolutionary conservation of the *MYEOV-*3’-enhancer and its interaction with *CCND1* in other species.

### 3.4. The *MYEOV*-3’-enhancer element in conserved in mammals and chicken

First, we questioned whether the *MYEOV*-3’-enhancer sequence was conserved in primates. We performed a BLAST search using the *MYEOV* locus as a query to look for homologous sequences in other primate species (**Supplementary Figure 5**). We discovered that not only was the coding sequence of *MYEOV* strongly conserved across all seven primate species considered, but its 3’ UTR sequence, where the human *MYEOV-*3’-enhancer is located, also shared this property (**Supplementary Figure 5**). Sequence conservation was not only seen in closely related species, such as chimpanzee and gorilla, but also in more distantly related species like squirrel monkeys (**Supplementary Figure 5**).

These observations suggest that both *MYEOV* and its 3’ UTR enhancer sequence are highly conserved outside of humans. To investigate if these regions also have enhancer activity in primates, we used the regulatory regions defined from histone a ChIP-seq dataset generated for lymphoblastoid cells of humans, chimpanzees, gorillas, orangutans and macaques (García-Pérez et al., 2021). We found that the entire *MYEOV*’s locus, including the homologous *MYEOV*-3’-enhancer region, is covered by strong enhancer or enhancer/promoter domains in the four non-human primates (**Figure 5**). This finding is particularly interesting since *MYEOV* is thought not to be transcribed in non-human primates (Papamichos et al., 2015), suggesting that this enhancer might be the only functional role in this region in these species. This is supported by data derived from orangutans, which are thought not to encode *MYEOV* due to a mutation leading to a premature stop codon (Papamichos et al., 2015), yet they still have an enhancer/promoter chromatin state present in the same region (**Figure 5**). This indicates that the *MYEOV* enhancer element might have predated the development of an ORF in this region.

**Figure 5.**
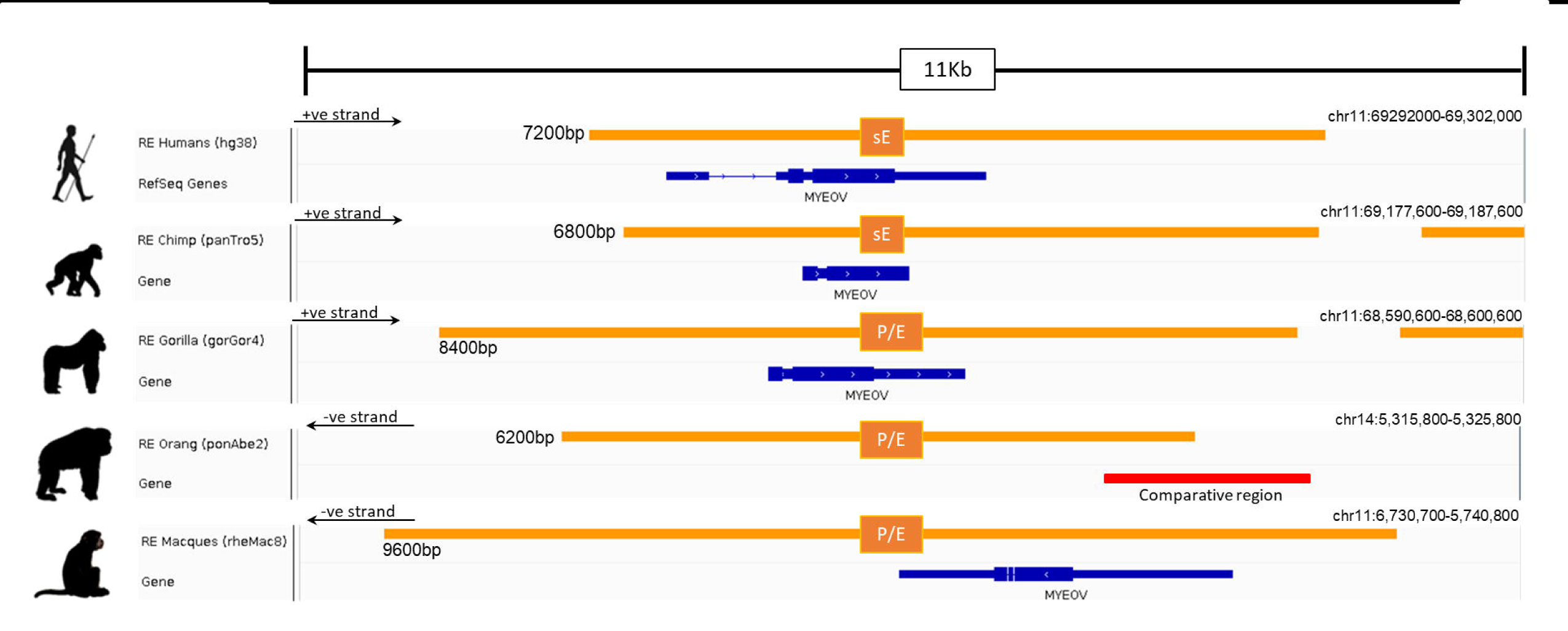
Regulatory state surrounding *MYEOV* in five primate species. Regulatory regions defined by ChromHMM in lymphoblastoid cells in human, chimpanzee, gorilla, orangutan, and macaque from García-Pérez et al. Abbreviation sE - Strong Enhancer, P/E - promoter/enhancer (where one replicate had an enhancer state and the other had a promoter state). Data visualised on IGV.

Next, we performed a comparative genomic analysis of additional species to determine if the *MYEOV* enhancer DNA sequence predated primates. BLAST searches across 17 mammalian species, and later expanded to include avian species, consistently observed BLAST hits ranging from 200-600 bp in a location similar to that between *CCND1* and *TPCN2* homologs across all species, barring rabbits (**Supplementary Figure 6**, **Supplementary table 4**) possibly due to the version of the rabbit genome being highly fragmented (Bai et al., 2021). To determine the conservation of this enhancer element, we performed multiple alignments across all these species using the DNA sequences surrounding these BLAST hits. Expanding the search out to fishes, such as zebrafish, no sequence homology was found but we identified a homologous region in chicken, again located between the chicken *CCND1* and *TPCN2* homologs (**Supplementary Figure 6**). We were able to show conservation across 18 different species, with a 273 bp sequence being evolutionarily conserved up to and including chickens (**Figure 6**).

**Figure 6.**
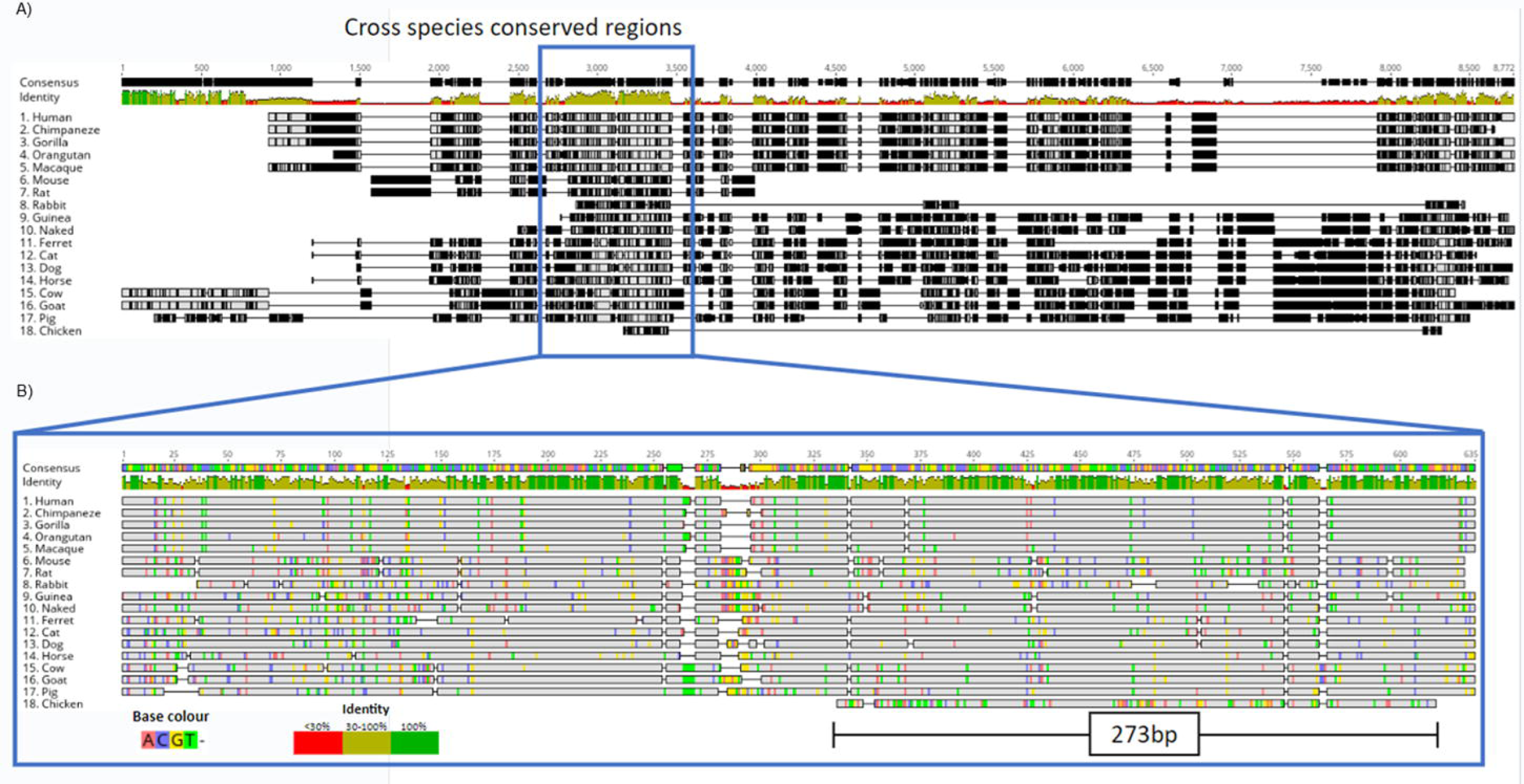
Multiple sequence alignment of BLAST hits reveals DNA sequence conservation of *MYOV*-3’-enhancer across mammals and chicken. A) Multiple sequence alignment illustrates the comparative analysis of a human *MYEOV*-3’-enhancer sequence with homologous sequences from 17 distinct species, identified through a BLAST search. The homologous sequences include 16 mammals, including 4 primates, and chicken. Sequence conservation is visualised using a colour-coded identity matrix, where darker colours signify higher sequence similarity. The consensus identity score is depicted, with yellow-grey colours indicating conservation ranging from 50% to 100%. **B)** This panel highlights the extraction and realignment of conserved regions, representing the largest contiguous block of identity scores between 50% and 100% present in all species. The realigned conserved region spans 635 bp for mammals and 273 bp for chicken. Identical residues are shaded in grey for clarity.

In some of these species there is publicly available ChIP-seq data of histone marks, so we checked the presence of marks compatible with enhancer activity. Using H3K27ac data across 20 mammalian species (Villar et al. 2015), we were able to observe H3K27ac marking this conserved region in mouse, rat, dog, bovine and porcine epigenomes (**Supplementary Figure 7**). Next, we used data ChIP-seq data for H3K4me1, H3K4me3, and H3K27ac from ten mammalian species (Roller et al., 2021), as well as another dataset including ChIP-seq data for the same histone marks in mice (Shen et al., 2012). Here, we found conservation of the enhancer state across macaques, mice, rats, dogs and cats with H3K4me1 and H3K27ac peaks surrounding the conserved region (**Figure 7**). Having observed sequence homology in chickens, we analysed ChIP-seq data for H3K4me3, H3K4me1 and H3K27ac derived from chicken liver tissue (Kern et al., 2021). H3K4me1 peaks were shown to surround the conserved region but no H3K27ac marks were detected, suggesting a potential poised enhancer state **(Figure 7**).

**Figure 7.**
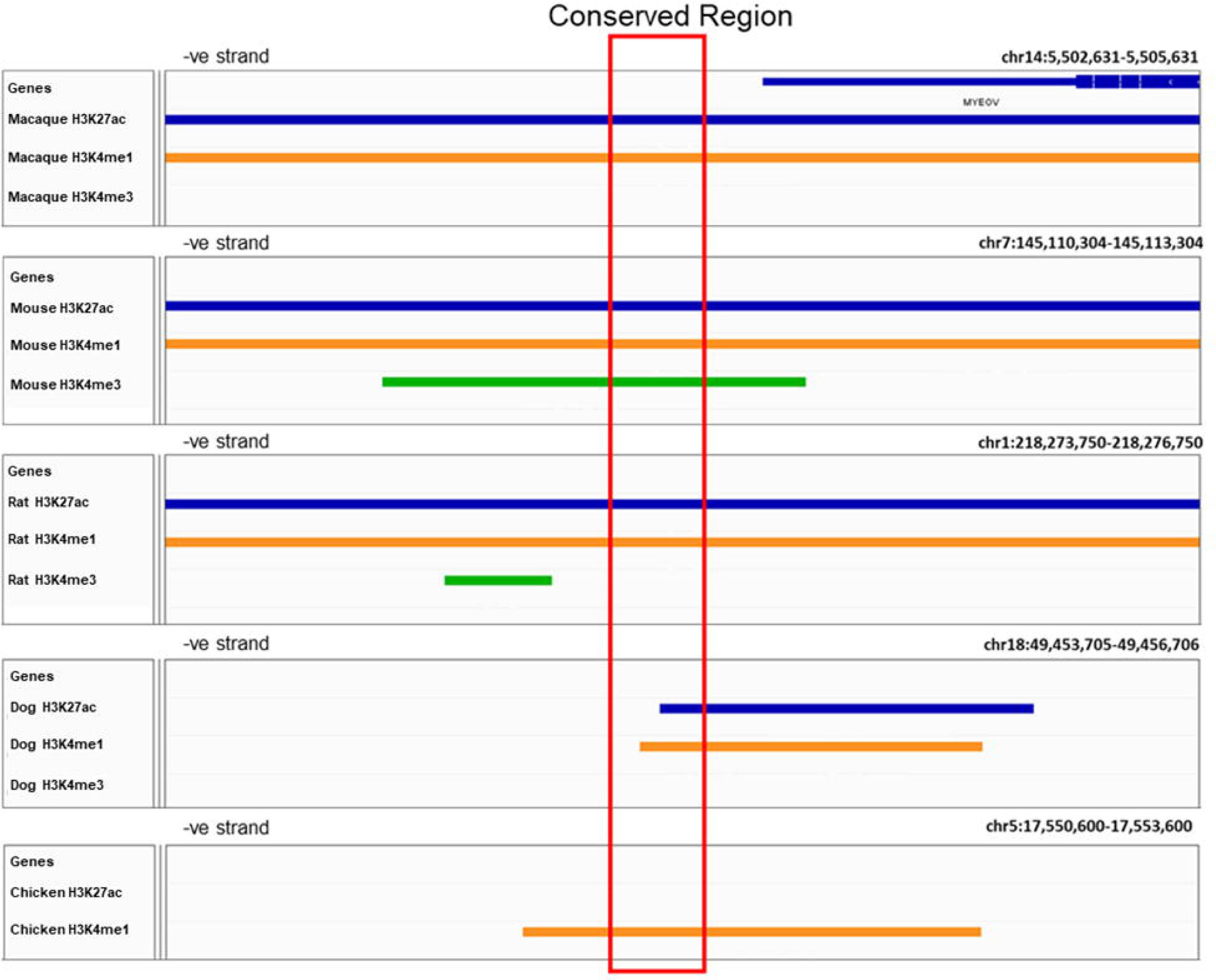
Conservation of enhancer chromatin state across multiple different species. ChIP-seq data for H3K4me1 (orange), H3K4me3 (green), H3K27ac (blue) derived from macaques, mice, rats, and dogs and chicken (Kern et al., 2021; Roller et al., 2021). Individual sample replicates combined into one track. The conserved region (the region with homologous sequence) is highlighted in the red box. Data visualised on IGV.

Taken together, these results show that the *MYEOV-*3’-enhancer is evolutionarily older than the *MYEOV’s* ORF and the patterns of histone marks in primates and non-primate mammals indicate a possible enhancer function of these homologous non-coding regions.

### 3.5. The mouse homolog of *MYEOV*-3’-enhancer interacts with *Ccnd1*

We have shown that the location of the homologs of *MYEOV-*3’-enhancer is located between the corresponding homologs of *CCND1* and *TCPN2* in 16 out of the 17 species analysed. This finding suggests that this conserved enhancer may have regulatory connections with the *CCND1* homologs in these species (as we observed in humans). To test this hypothesis, we characterised the mouse homolog of *MYEOV*-3’-enhancer (m*MYEOV*-like-enhancer) in more detail.

We analysed data from Mouse ENCODE for four different murine cell/tissue types to determine if this enhancer state was present in tissues other than the liver. We analysed H3K4me1 and H3K27ac data from CH12 cells (a lymphoma cell line used as a B cell model), lung tissue, liver tissue and bone marrow (**Supplementary Figure 9**). Similar to the equivalent human tissues, H3K4me1 and H3K27ac peaks covered the conserved region in all of these cell types (**Supplementary Figure 9**). We also noticed that upstream of the m*MYEOV*-like-enhancer, there is a CTCF site to which CTCF had bound in B cells (**Supplementary Figure 9**). This suggested that this enhancer might also be targeting the *Ccnd1* in mice, which is located in a similar location as seen in humans (**Supplementary Figure 9**).

To investigate whether the m*MYEOV*-like-enhancer also targets *Ccnd1* in mice, we analysed PCHi-C data performed in both murine pre-B cells (Koohy et al., 2018) and mESCs (Schoenfelder et al., 2015) as well as DNase capture Hi-C data from mESCs (Joshi et al., 2015). We observed that in pre-B cells of old mice, *CCND1* interacts with this conserved enhancer region (**Figure 8A-B**). This interaction can be also found in DNase capture Hi-C data performed in mESCs, which highlights interactions between open chromatin regions (**Figure 8C**). However, in PCHi-C data from mESCs (Schoenfelder et al., 2015), no interaction is seen with the evolutionarily conserved region, but an interaction is seen with a region just downstream of this element (**Figure 8D**). The difference in mESC capture data might be due to differences in the regions captured, as the *CCND1* bait is located at chr7:152,118,105-152,126,754 whereas in DNase capture Hi-C, the *CCND1* interacting site is at chr7:152,121,377-152,134,010.

**Figure 8.**
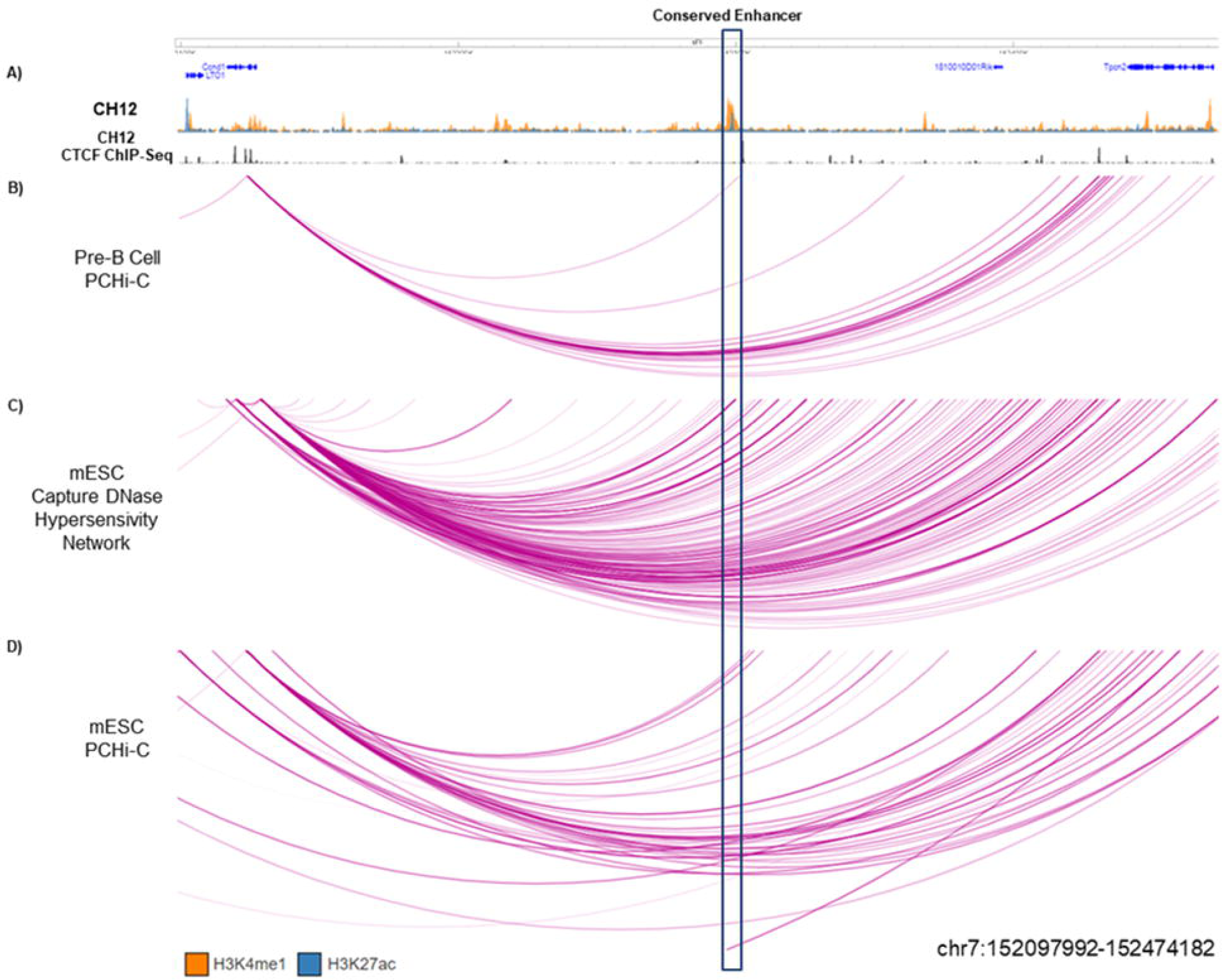
Interaction observed between conserved enhancer sequence and *CCND1*. **A)** Top first panel shows ChIP-seq data for H3K4me1 (Orange), H3K27ac (Blue), second panel shows CTCF in CH12 mouse lymphoma cell line. **B)** PCHi-C performed in pre-B cells in old mice (Koohy et al., 2018). **C)** DNase capture Hi-C data from mESC (Joshi et al., 2015). **D)** PCHi-C performed in mESC (Schoenfelder et al., 2015) The purple arcs denote interactions between fragments. The conserved region (the region with homologous sequence) is highlighted in the blue box. Data visualised using the WashU Epigenome Browser.

In conclusion, these experimental data reveal that the genomic region containing the m*MYEOV*-like-enhancer shows chromatin interactions with *Ccnd1* in mice, mirroring the interactions we observed between the *MYEOV*-3’-enhancer and *CCND1* in humans.

## 4. Discussion

An active enhancer in B cells situated at the 3’ end of *MYEOV*, which we call the *MYEOV*-3’-enhancer, has recently been reported (Mikulasova et al., 2022). In this study we found that this enhancer exhibited activity across a diverse array of both healthy and disease-afflicted human cell types, thereby underscoring its pivotal role in regulating gene expression within varied contexts.

Although the exact protein function of MYEOV remains elusive, ongoing studies highlight potential non-coding oncogenic functions at the transcript level. Overexpression of *MYEOV* has been associated with poor prognosis in different cancers (Fang et al., 2019; Liang et al., 2020); (Moreaux et al., 2010); (Janssen et al., 2002a); (Janssen et al., 2002b); (Leyden et al., 2006); (Moss et al., 2006); (Takita et al., 2011); (Shen et al., 2021). Notably, transcripts of *MYEOV* possess oncogenic properties due to their capability to sequester miRNAs, thus preventing these molecules from targeting and repressing the expression of oncogenic factors (Fang et al., 2019).

Through an analysis of enhancer-associated histone marks (H3K4me1, H3K4me3, and H3K27ac) in the syntenic region, we unveiled a striking conservation of the enhancer chromatin state across mammals. The *MYEOV*’s ORF is a young gene, with an ORF exclusive to primates but the DNA sequence of the core enhancer region exhibited conservation not only across mammals but also in chickens. Our results indicate that the histone marks associated with enhancer activity preceded the emergence of the ORF of *MYEOV*.

Sequence homologs of this novel *MYEOV*-3’-enhancer were discovered through BLAST searches, where it was consistently situated adjacent to the homologs of *CCND1*. The synteny observed between *MYEOV* and *CCND1* across various species strongly suggests a functional relationship between these two elements. Furthermore, our research revealed that in human and murine cells (which lacks an ORF, yet bear the enhancer marks) the *MYEOV*-3’-enhancer and *CCND1/Ccnd1* display interactions within the 3D genome. This suggests deep conservation (Wong et al., 2020) of this enhancer that extends beyond the ORF shared by most primates

The *MYEOV*-3’-enhancer shows activity across a broad spectrum of human tissues, as indicated by the presence of H3K27ac. Consequently, our interpretation posits that the *MYEOV*-3’-enhancer contributes to the regulation of *CCND1* expression. In non-pathogenic cells, the expression of *CCND1* is likely under the regulation of polycomb repressive complexes that exert their influence on its promoter region (Mikulasova et al., 2022; Rico et al., 2022). While the *MYEOV* gene is active and spatially proximate to *CCND1*, the expression of *CCND1* hinges on the removal of polycomb-mediated inhibition. Conversely, in normal cells, the transcription of *MYEOV* remains inactive, largely attributed to DNA methylation (Fang et al., 2019; Liang et al., 2020). *CCND1* seems to have an association with more than one enhancer, as supported by the observation of multiple 3D genome interactions with regions bearing enhancer histone marks that extend beyond those linked with *MYEOV* (Rico et al., 2022). Future enhancer deletion/perturbation experiments using CRISPR (Kent et al., 2023) will help to prove whether the observed interactions between *CCND1* and *MYEOV*-3’-enhancer are indeed regulatory.

In the context of cancer, the insertion of super-enhancer elements from different chromosomes into the intergenic region flanked by the *MYEOV*-3’-enhancer and *CCND1* culminates in the emergence of an expansive active regulatory domain spanning the *CCND1* gene locus. This occurrence results in the displacement of repressive polycomb-associated signatures, ultimately paving the way for the activation of *CCND1* expression (Mikulasova et al., 2022). Intriguingly, this scenario concurrently activates the *MYEOV*’s gene body, as evidenced by the appearance of active promoter-associated marks (an H3K4me3 broad domain) over the *MYEOV*’s gene body (Mikulasova et al., 2022). This suggests that the introduction of these super-enhancer elements not only upregulates *CCND1* expression but also augments the transcriptional activity of *MYEOV* by creating a conducive chromatin environment through the acquisition of active promoter marks (Kent et al., 2023).

In cancer, documented cases of DNA amplification at 11q13, primarily likely selecting for *CCND1* proto-oncogene activation, often correspond to elevated *MYEOV* expression and an unfavourable prognosis (Fang et al., 2019; Liang et al., 2020). We speculate that during amplification events, duplicated genomic segments undergo the loss of their repressive epigenomic signatures, consequently enabling demethylation of *MYEOV*, and thereby leading to an overexpression of the *MYEOV*’s RNA in cancer cells, and/or permitting the *MYEOV-3’-* enhancer to regulate *CCND1*, thereby boosting *CCND1* overexpression.

The concept of chromatin accessibility and enhancer characteristics that facilitate the birth of new genes has been proposed in prior research (Majic and Payne, 2020). Our findings align with this concept, effectively illustrating the potential of enhancers as a genomic environment that facilitates the emergence of new genes. Beyond its role in regulating *CCND1*, the *MYEOV*-3’-enhancer’s significance in oncogenesis becomes all the more pronounced. It might actively contribute to oncogenic processes through DNA amplifications or chromosomal translocations, potentially allowing it to effectively regulate the expression of other proto-oncogenes, thereby expanding its impact on the dynamics of cellular function.

Finally, we would like to highlight that the conservation of the *MYEOV*-3’-enhancer element across species provides a unique opportunity for the development of animal models, such as mouse or chicken, aimed at studying enhancer function and its interactions. While the creation of knockdown models of *MYEOV* in mice is impossible due to the gene’s absence, the preservation of the enhancer elements offers a promising avenue for functional exploration of the roles of *MYEOV*-3’-enhancer in carcinogenesis.

## Supporting information

Supplementary Tables (one per tab)

Supplementary Table and Figures Descriptions

## 7. Supplementary figure legends

**Supplementary Figure 1: NIH Roadmap data highlighting chromatin state surrounding MYEOV’s Locus.** H3K4me1 (Orange), H3K4me3 (Green) and H3K27ac (Blue) ChIP-seq data taken from NIH Roadmap data for GM12878, B cells, Lung, Liver and Pancreatic Tissues alongside GM12878 and B cell DNase-seq. Data was visualised on WASHU genome browser.

**Supplementary Figure 2: Interaction observed when CCND1 is taken as viewpoint**. Promoter capture Hi-C data across 5 different tissues/cell line taken from Jung et al.; these being GM12878/GM19240 Lymphoblastoid Cell line, Lung, Liver and Pancreatic tissues and H1-derived Mesenchymal Stem cells. CCND1 was taken as bait and interactions were filtered to only show significant interaction over −log10(P-value) = 2 and to only show normalised count. Data was visualised on 3DIV.

**Supplementary Figure 3: Interaction observed when MYEOV is taken as the viewpoint.** Promoter capture Hi-C data across 5 different tissues/cell line taken from Jung et al.; these being GM12878/GM19240 Lymphoblastoid Cell line, Lung, Liver and Pancreatic tissues and H1-derived Mesenchymal Stem cells. MYEOV was taken as bait and interactions were filtered to only show significant interaction over −log10(P-value) = 2 and to only show normalised count. Data was visualised on 3DIV.

**Supplementary Figure 4: Interaction observed when LTO1/ORAOV1 is taken as bait.** Promoter capture Hi-C data across 5 different tissues/cell line taken from Jung et al.; these being GM12878/GM19240 Lymphoblastoid Cell line, Lung, Liver and Pancreatic tissues and H1-derived Mesenchymal Stem cells. ORAOV1 (LTO1) was taken as bait and interactions were filtered to only show significant interaction over −log10(P-value) = 2 and to only show normalised count. Data was visualised on 3DIV.

**Supplementary Figure 5: BLAST Hit for MYEOV locus seen in 7 primate species.** Ensembl BLAST search was performed using MYEOV’s locus as a query against 7 primate species which included chimpanzee (*Pan troglodytes*), Gorilla (*Gorilla gorilla*), Orangutan (*Pongo abelii*), Gibbon (*Nomascus leucogenys*), Rhesus Macaque (*Macaca mulatta*), Marmoset (*Callithrix jacchus*), Squirrel Monkey (*Saimiri boliviensis*). Conserved sequences are shown as red rectangles with the percentage ID of blast hit determined by the shade. Data was visualised on the Ensembl.

**Supplementary Figure 6: Conservation of Gene synteny surrounding BLAST sequence.** Region which surrounds Ensembl BLAST top hit was located across 5 different species, these being Human, Dog, Cow, Goat and Chicken when MYEOV 3’UTR was used as the query sequence. Each BLAST hit is denoted with a red line. Data was visualised on the Ensembl.

**Supplementary Figure 7: Conservation of H3K27ac signal across Mammalian Tissue**. H3K27ac (Blue) and H3K4me3 (Orange) ChIP-seq data taken from liver tissue of 6 species including Humans, Mouse, Rat, Dog, Cow and Pig (Villar et al. 2015). Also, H3K4me1 and H3K4me3 ChIP-seq data from mouse liver tissue (Shen et al. 2012). Conserved region highlighted in red is the homologous sequence determined from multiple sequence alignment. Data was visualised on IGV.

**Supplementary Figure 8: Comparing Histone data from Roller et al. to Villar et al**. ChIP-seq data for 3 mammalian species was taken from both Roller et al and Villar et al. This included H3K27ac (Blue), H3K4me1 (Orange) and H3K4me3 (Green) for Rat, Dog and Pig Liver tissue. Conserved region highlighted in red is the homologous sequence determined from multiple sequence alignment. Data was visualised on IGV.

**Supplementary Figure 9: Conservation of enhancer chromatin state across multiple mouse tissues**. Top Panel: Highlights ChIP-seq for H3K4me1 (Orange), H3K4me3 (Green) and H3K27ac (Blue) in CH12 mouse lymphoma cell line, lung, liver and bone marrow tissue in mice. Alongside, ChIP-seq data for CTCF in CH12 mouse lymphoma cell line. Bottom Panel: Show zoom in region surrounding conserved region which is highlighted in yellow and is the homologous sequence determined from multiple sequence alignment.

