## Supplementary Table and Figures Descriptions for "The discovery of an evolutionarily conserved enhancer within the MYEOV locus suggests an unexpected role for this non-coding region in cancer"

**Legends of Supplementary Tables**

**Supplementary Table 1.** This supplementary table presents sources used for studying chromatin states and regulatory regions across different species and tissues. It contains sources of chromatin profiling data, including ChIP-seq and ATAC-seq, for multiple species and tissues. The data include information on histone marks (H3K4me1, H3K4me3, H3K27ac, H3K27me3, H3K36me3) and are sourced from diverse studies involving primates, other mammalian species, and chicken.

**Supplementary Table 2**. This supplementary table presents the results of an extensive analysis of the presence or absence of H3K27ac peaks within MYEOV's 3'UTR region in a wide range of human tissues, cell lines, and cell types, shedding light on the tissues where this enhancer is active.

**Supplementary Table 3.** This supplementary table presents the data resources used to investigate the regulatory roles of MYEOV's enhancer element in both human and mouse genomes and identification of gene targets of MYEOV's enhancer. The table includes sources of ChIP-seq data for histone modifications (H3K4me1, H3K4me3, H3K27ac) and CTCF in multiple mouse tissues, including CH12 mouse B cell lymphoma cell line, adult liver cells, lung liver cells, and bone marrow. Additionally, it provides sources of human PCHi-C studies in specific tissues associated with cancer. The table also contains data on long-range chromatin interactions, ChIA-PET data, and CTCF data to explore potential mechanisms of enhancer-target promoter interactions and confirm topologically associated domain (TAD) structures.

**Supplementary Table 4.** This supplementary table presents DNA sequence conserved regions of MYEOV's enhancer element across a diverse set of 17 species, pinpointed by BLASTN search, and multiple sequence alignments, are documented in this table.
