## Supplementary figures and images for "The discovery of an evolutionarily conserved enhancer within the MYEOV locus suggests an unexpected role for this non-coding region in cancer"

### Supplementary Figure 1

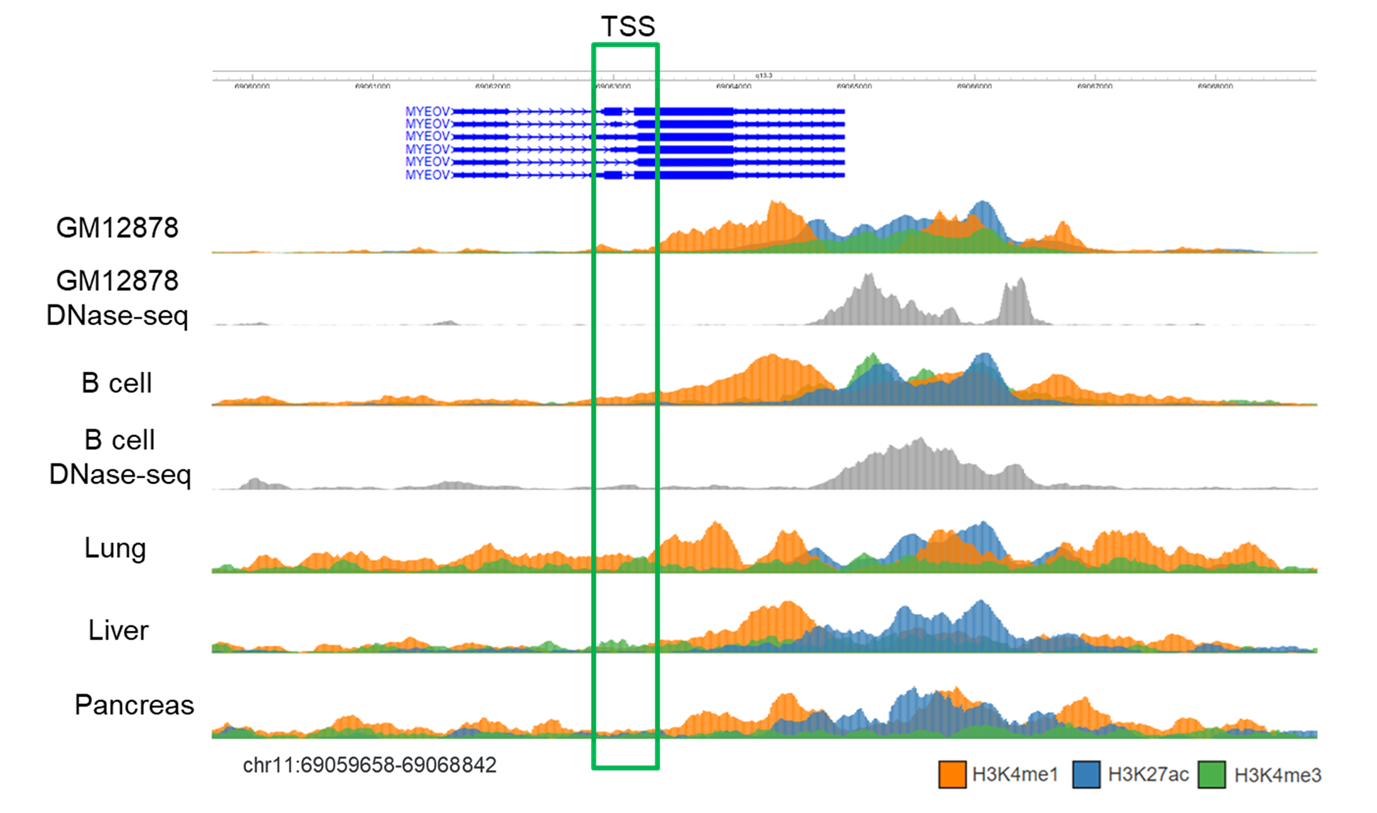

### Supplementary Figure 2

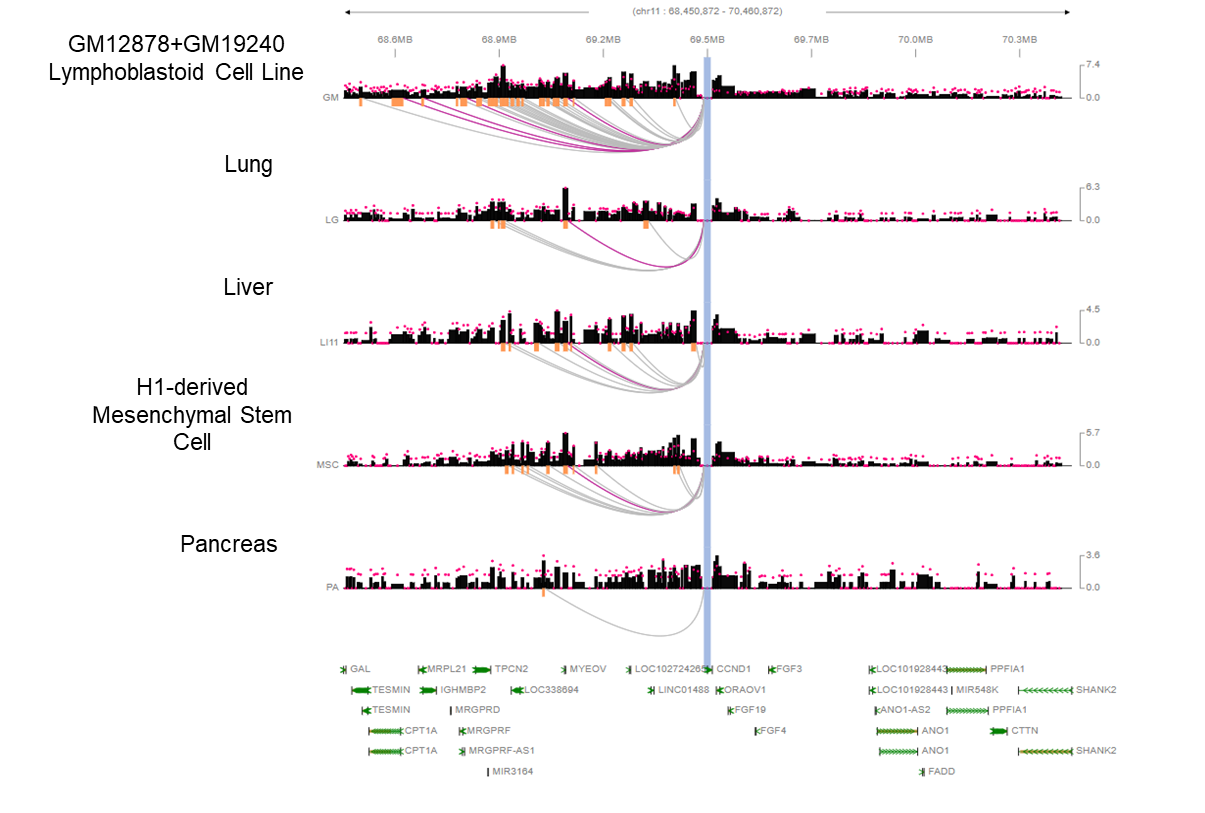

### Supplementary Figure 3

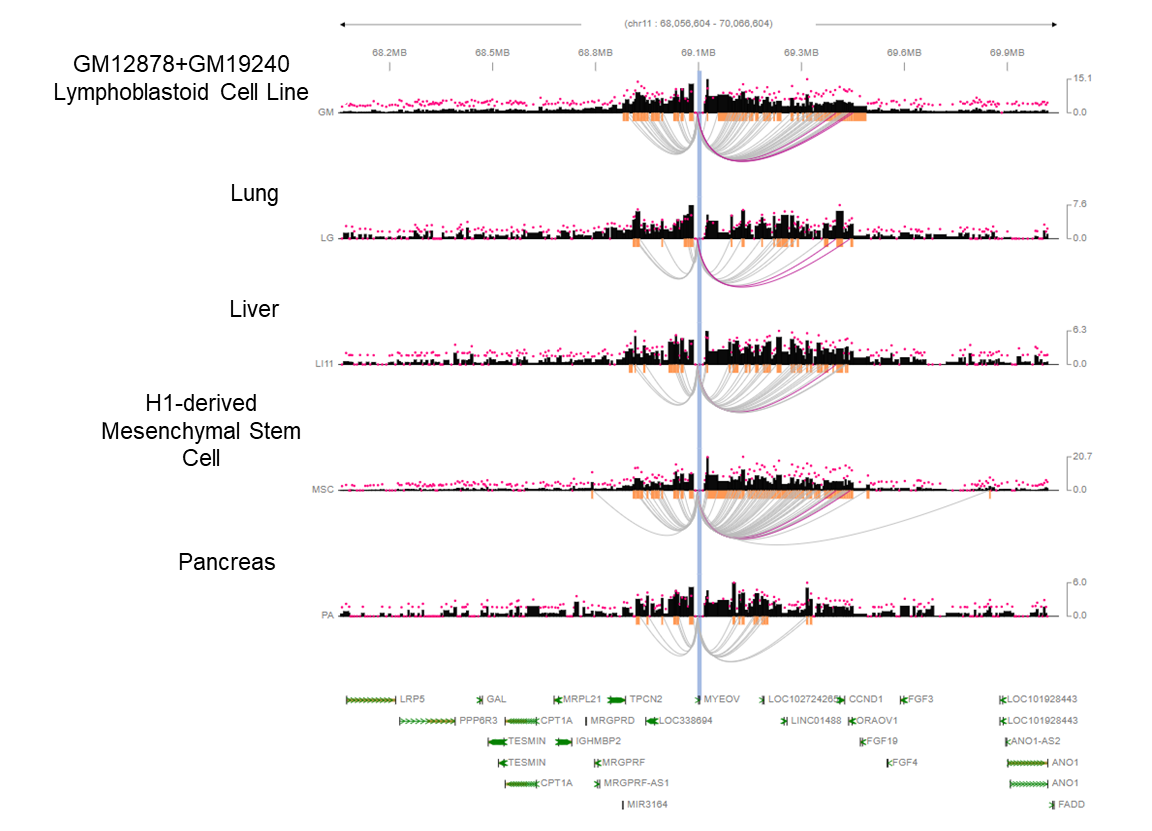

### Supplementary Figure 4

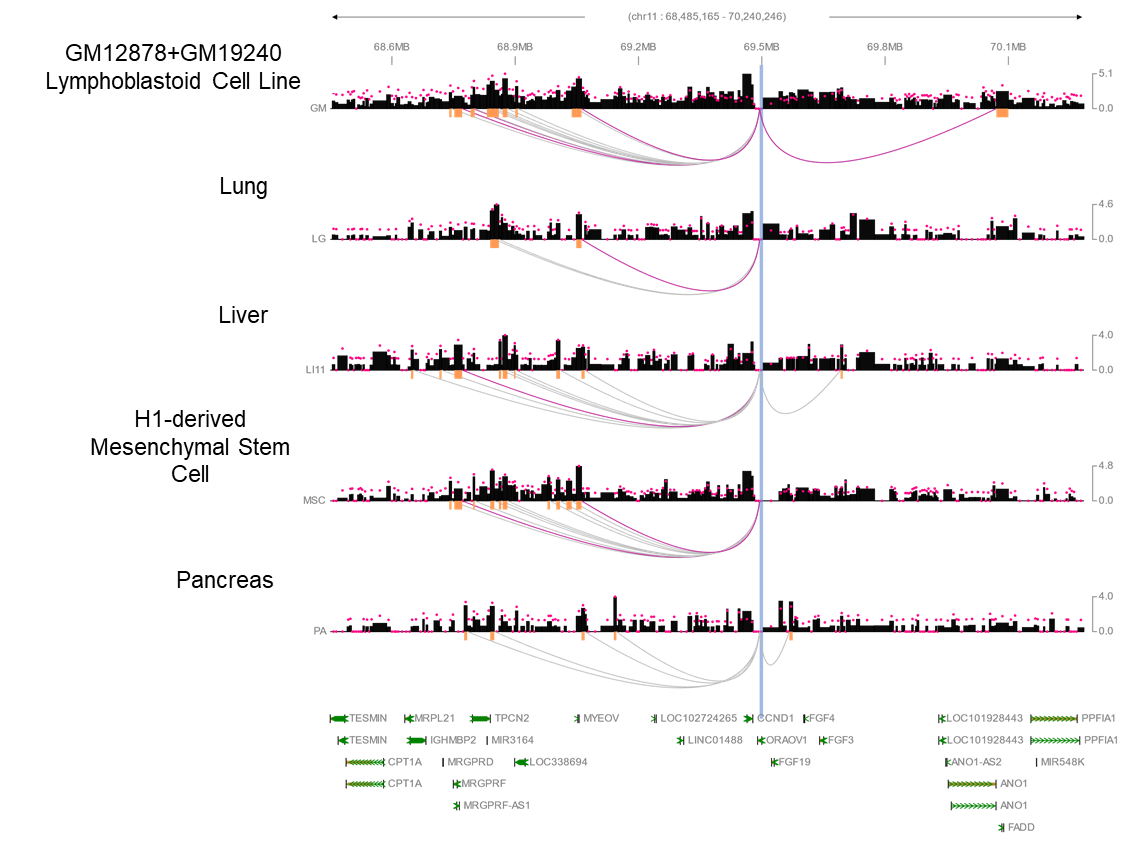

### Supplementary Figure 5

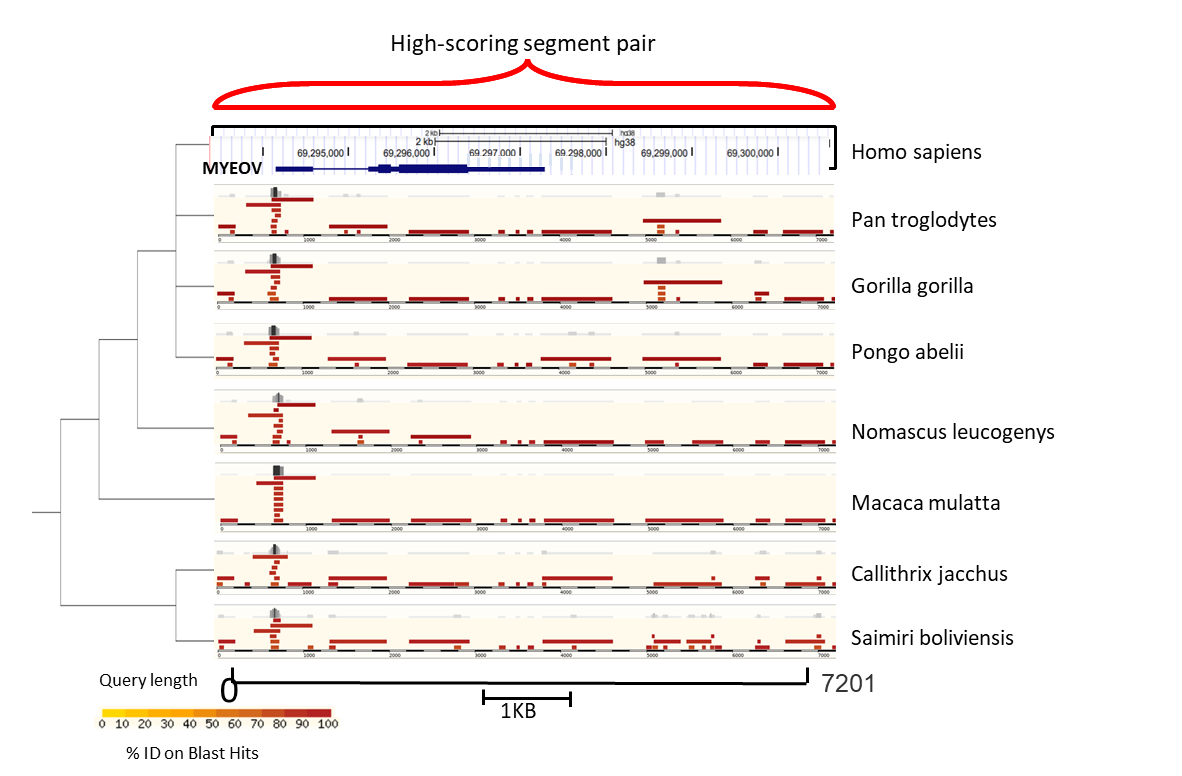

### Supplementary Figure 6

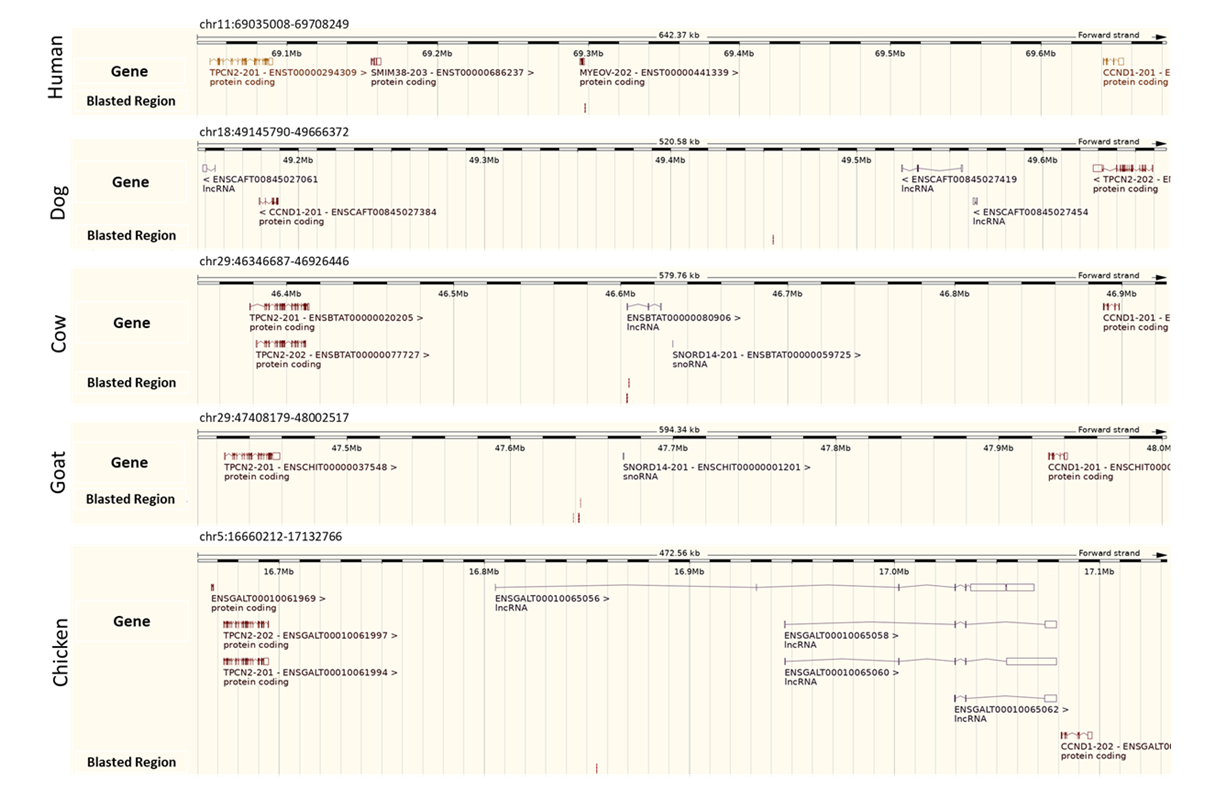

### Supplementary Figure 7

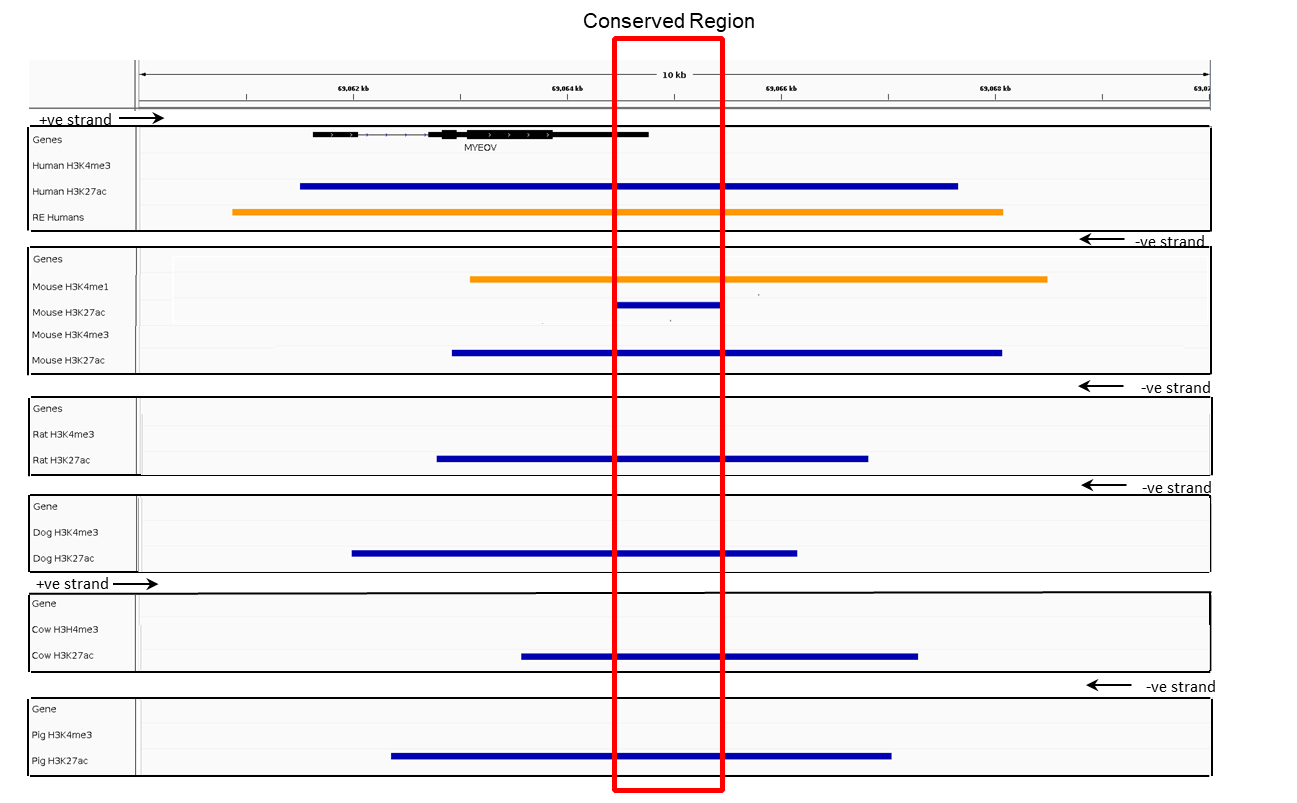

### Supplementary Figure 8

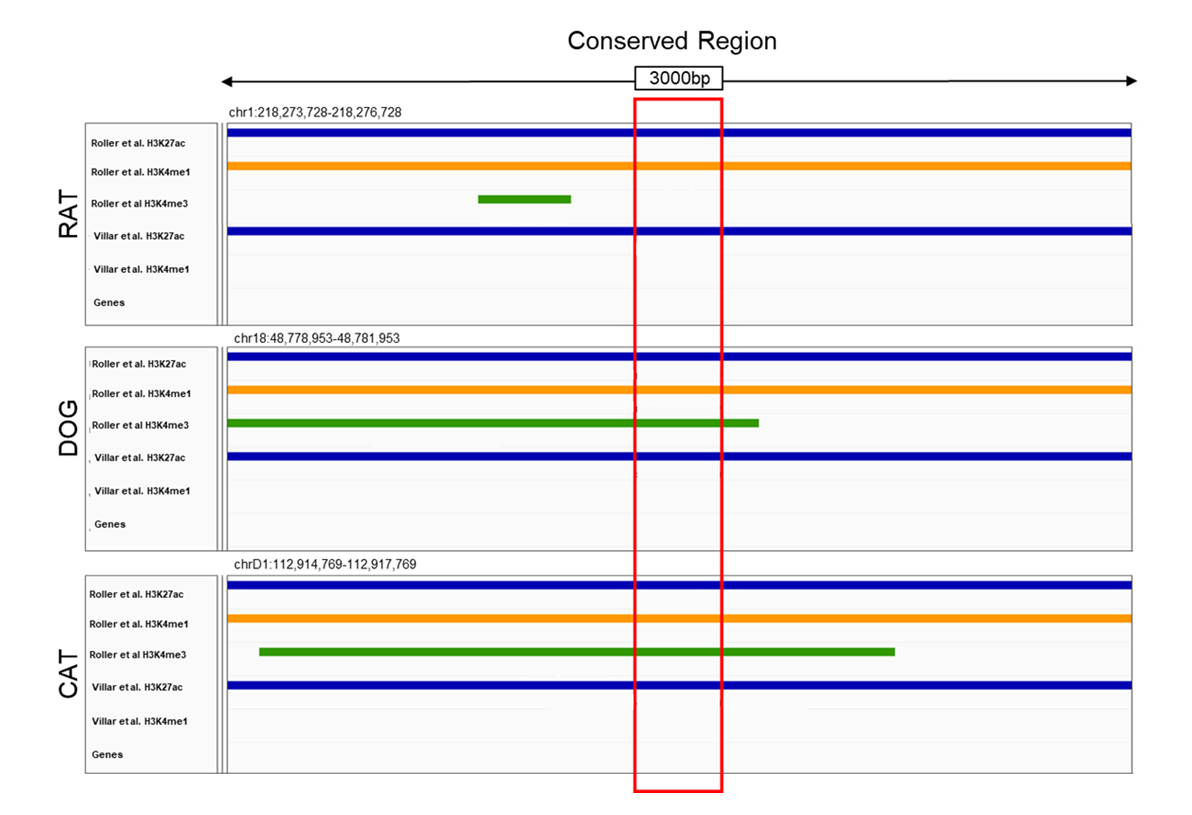

### Supplementary Figure 9

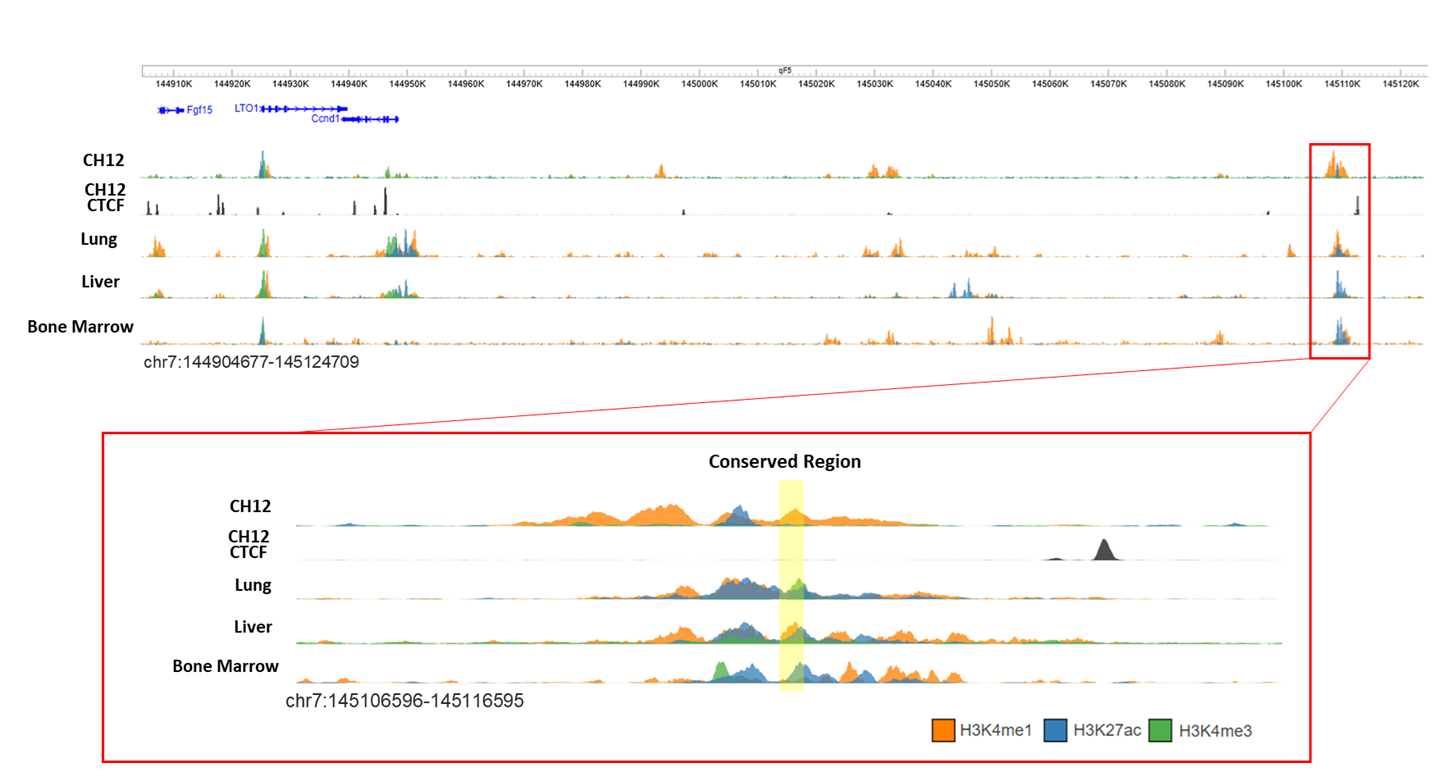
